# Dimerization of Cdc13 is essential for dynamic DNA exchange on telomeric DNA

**DOI:** 10.1101/2025.03.25.645294

**Authors:** David G. Nickens, Spencer J. Gray, Robert H. Simmons, Matthew L. Bochman

## Abstract

Single-stranded DNA (ssDNA) binding proteins (ssBPs) are essential in eukaryotes to protect telomeres from nuclease activity. In *Saccharomyces cerevisiae*, the ssBP Cdc13 is an essential protein that acts as a central regulator of telomere length homeostasis and chromosome end protection, both alone and as part of the Cdc13-Stn1-Ten1 (CST) complex. Cdc13 has high binding affinity for telomeric ssDNA, with a very slow off-rate. Previously, we reported that despite this tight ssDNA binding, Cdc13 rapidly exchanges between bound and unbound telomeric ssDNA substrates, even at sub-stoichiometric concentrations of competitor ssDNA. This dynamic DNA exchange (DDE) is dependent on the presence and length of telomeric repeat sequence ssDNA and requires both Cdc13 DNA binding domains, OB1 and OB3. Here we investigated if Cdc13 dimerization is important for DDE by characterizing the dimerization mutant Cdc13-L91R. Using mass photometry, we confirmed that Cdc13-L91R fails to dimerize in solution, even in the presence of ssDNA. Gel-based DDE assays revealed that Cdc13-L91R fails to undergo ssDNA exchange compared to recombinant wild-type protein. Biolayer interferometry demonstrated that this effect was not due to differences in ssDNA binding kinetics. Thus, dimerization of Cdc13 is essential for DDE, and we model how this may impact telomere biology *in vivo*.

**GRAPHICAL ABSTRACT:** 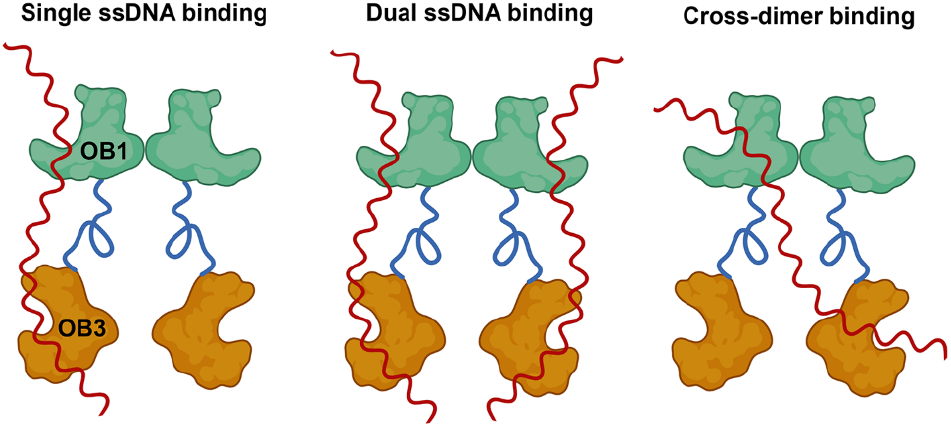

## INTRODUCTION

Telomeres are essential nucleoprotein structures involved in nuclease protection of linear chromosome ends to prevent chromosomal fusion or recombination of telomeric 3□ ssDNA overhangs (1-5). In *Saccharomyces cerevisiae*, an important component of telomere end protection is the CST complex, a heterotrimer formed by Cdc13, Stn1, and Ten1 (5-7). Cdc13 is the primary ssDNA binding protein (ssBP) of the CST complex, and it binds with high affinity to G-rich DNA, giving Cdc13 specificity for yeast telomeres with the TG_1-3_ repeat sequence (8-10). In addition to its end protection functions, Cdc13 also plays several important regulatory roles in control of telomere length homeostasis (11-14). These include the dissociation of the CST complex leading to activation of telomerase, the interaction with the catalytic subunit of Pol α-primase, and the reassociation of CST following telomere elongation (15-17). Therefore, Cdc13 efficiently binds to G-rich ssDNA and acts as a central regulator of DNA end protection and telomerase activity in budding yeast (8,18,19).

The Cdc13 polypeptide has a complex organization of DNA binding domains and protein interfaces for interactions with DNA polymerase α, the Est1 subunit of telomerase, the nuclear import machinery, Stn1 and Ten1, and homodimerization domains (20,21). Four oligonucleotide/oligosaccharide-binding (OB) domains have been identified in Cdc13 (20,22). These include OB1, the low-affinity DNA binding domain, and OB3, which contains a high-affinity DNA binding interface (8,23). Interestingly, the structure of OB1 is very similar to that of OB3, though OB1 lacks the long loop region reported in OB3 (22,24,25). The OB1 domain also contains a hydrophobic region that is the primary dimerization interface for Cdc13 (22).

The OB2 and OB4 domains do not have characterized *in vitro* DNA binding activity. Although a temperature-sensitive mutation in OB2 causes a loss of DNA binding at non-permissive temperatures, leading to the rapid export of Cdc13 from the nucleus (20), the effect on DNA binding is thought to be indirect. The Stn1 and Ten1 binding interfaces are located within the OB2 and OB4 folds (26), and the telomerase recruitment domain and the nuclear localization domain lie between the OB1 and OB2 domains (20). The OB2 and OB4 domains have both been studied in isolation, and both form homodimers (13,16). Mutations in OB2 are reported to disrupt Stn1 binding, confirming that an interaction between OB2 and OB4 is essential for CST complex formation (13).

The role of Cdc13 as a central regulator of telomere stability requires that it is bound at or near 3□ telomeric ssDNA to protect against nuclease activity (18,27-29). During G1 phase in *S. cerevisiae*, the CST complex is bound to short 3□ telomeric ssDNA overhangs of 12-18 nt (17). As cells transition from G1 phase and enter S phase, phosphorylation of Cdc13 and Stn1 lead to dissociation of the CST complex, while Cdc13 remains ssDNA bound (15,30,31). Following this dissociation event, Cdc13 association with Stn1 is replaced by interactions with the Est1 component of telomerase, which activates the Est2 reverse transcriptase, leading to telomerase extension (15,32,33). As telomere elongation proceeds, Cdc13 becomes hyper-phosphorylated and SUMOylated, while Stn1 is dephosphorylated, leading to a reassociation of the CST complex (17,34). As this is occurring, movement of the replication complex and control of DNA Pol α-primase activity requires that Cdc13 is removed and relocated to the newly synthesized telomere 3□ end to reform the CST complex (16). Due to the tight binding of the Cdc13 OB3 to telomeric ssDNA, a significant amount of energy would be required for Cdc13 to dissociate from its pre-replication binding site (23). An additional complication of such a dissociation is that unbound Cdc13 is rapidly exported from the nucleus (20). This raises the question, how is Cdc13, a ssBP with high affinity for telomeric repeat sequence DNA, moved from one position to another on its ssDNA substrate to complete replication and restore 3□ end protection?

Several groups, including our own, have investigated if the Pif1 helicase can remove Cdc13 from telomeric ssDNA *in vitro* (9,35,36), but the conclusions are mixed, with evidence presented that Pif1 both can and cannot remove Cdc13 from ssDNA depending upon the biochemical conditions used. Important variables for Cdc13 removal from ssDNA by Pif1 include the length of ssDNA that Cdc13 is bound to, the presence of available ssDNA for Pif1 loading, and the condensation state of the Cdc13-ssDNA complex (35,36). A recent report used helicase assays to monitor the removal of Cdc13 from duplex DNA substrates with ssDNA tails, allowing Pif1 to unwind the duplex region (36). In this work, as the length of the ssDNA increases, the rate of Cdc13 removal by Pif1 also increases due to tandem helicase activity. Other groups have reported that Pif1 inhibits telomerase *in vivo* at all telomeres, regardless of the length of the ssDNA overhang, and possibly independent of Cdc13 (37). Telomere DNA length has been proposed as an important variable for control of Cdc13 removal by Pif1 (35,38). Single-molecule analysis with immobilized telomere mimics indicates that Cdc13 condenses the DNA, forming a higher-order structure that blocks Pif1 from displacing Cdc13 (35).

These authors also report that Pif1 can remove Cdc13 from a ssDNA telomere mimic. Reports using HO endonuclease-generated double strand breaks (DSBs) indicate that long telomere constructs (34-82 bp) are insensitive to Pif1 activity, allowing increased telomerase extension (38,39). It is hypothesized that this represents a transition point between DSBs, represented by (TG_1-3_)_n_ dsDNA > 34 bp, and short telomeres, represented by (TG_1-3_)_n_ dsDNA < 34 bp. The authors propose that Pif1 cannot remove Cdc13 from the short telomere mimics, allowing telomerase extension to proceed (38). While it is hard to draw firm conclusions from such conflicting data, it is clear that Pif1 can remove Cdc13 from telomeric ssDNA under certain conditions.

However, we recently found that at physiological temperatures, Cdc13 itself rapidly exchanges between bound and unbound telomeric ssDNA substrates in a process that we termed dynamic DNA exchange (DDE) (9). Indeed, DDE is significantly faster than Cdc13 eviction from ssDNA by Pif1 or the combination of the Pif1 and Hrq1 helicases (9,36). DDE by Cdc13 is a temperature-dependent phenomenon that does not require ATP, Mg^++^, or additional proteins. Efficient exchange between telomeric repeat sequence ssDNAs occurs at physiological temperatures, but no DDE is observed at 4°C. Concerning the nucleic acid, we demonstrated that both the length and sequence of the unbound secondary ssDNA affect DDE. Increasing the length of the secondary substrate from 11 to 50 nt proportionately increases the extent of Cdc13 exchange from a prebound 30-nt primary substrate. Increasing the length of the primary substrate only weakly correlated with DDE activity, but once the length of prebound ssDNA reaches 50 nt, DDE activity is significantly reduced. Further, DDE is only observed with telomeric sequence ssDNAs; poly(dT) ssDNA substrates of equal length to the telomeric substrates do not support DDE. Thus, DDE by Cdc13 also demonstrates physiological sequence specificity. We hypothesized that reduced DDE activity as longer telomeric ssDNA is boundby Cdc13 could be part of a mechanism for controlling telomere length homeostasis *in vivo*.

Using truncation mutants of Cdc13, we were also able to interrogate the protein determinants of DDE. We found that DDE requires both the high-(OB3) and low-affinity (OB1) DNA binding domains (9). Recombinant OB3 domain in isolation tightly binds to telomeric ssDNA, even in the presence of a large molar excess of competitor secondary substrate. Similarly, an N-terminal truncation mutant that includes OB3 but lacks the OB1 domain (Cdc13ΔOB1) also lacks DDE activity. The OB1 domain in isolation displays low-affinity ssDNA binding, as previously reported (22,40), complicating measurement and comparison of DDE (9). However, biolayer interferometry (BLI) analyses (41) suggest that competition for primary *vs*. secondary ssDNA substrate binding by OB1 is likely driven by its high on-and off-rates rather than DDE (9). We concluded that DDE by Cdc13 requires both the OB1 and OB3 domains, but because Cdc13 exists as a homodimer in solution (22), it is unclear if DDE requires the OB1 and OB3 domains from each subunit or from a single monomer within the dimer. Here, we used mass photometry, DDE assays, BLI, and three-dimensional structure prediction to confirm that the Cdc13-L91R mutant is a ssDNA binding-competent monomer in solution and compare its ability to undergo DDE to wild-type dimeric Cdc13. We propose an updated model for how Cdc13 moves to the new telomere end following telomerase extension, where DNA binding at alternate open DNA binding domains induces significant conformational changes that are communicated across the dimer interface, leading to the release of Cdc13 and DDE-based movement along ssDNA without dissociation.

## MATERIALS AND METHODS

### Reagents

All oligonucleotide substrates used in this work were purchased from Integrated DNA Technologies (Coralville, IA, USA) and included: IR700-Tel30G (/5IRD700/CGCCATGCTGATCCGTGTGGTGTGTGTGGG), IR800-Tel30G (/5IRD800/CGCCATGCTGATCCGTGTGGTGTGTGTGGG), biotinylated Tel30G (/5BiotiTEG/CGCCATGCTGATCCGTGTGGTGTGTGTGGG), and unlabelled Tel30G and Tel50G (GTGTGGGTGTGGTGTGGGTGTGGTGTGGGTGTGTGGGTGTGGTGTGGGTG). All other chemical reagents were of molecular biology grade or higher and purchased from Sigma-Aldrich (St. Louis, MO, USA), unless otherwise noted.

### Biological resources

Vectors for over-expression of full-length, wild-type Cdc13 and Cdc13-L91R in *Spodoptera frugiperda* Sf9 tissue culture were provided by Dr. Hengyao Niu. These pFastBac-based vectors (Invitrogen, Waltham, MA, USA) encode a 5□ -SUMO-His_6_ tag and a 3□ -FLAG tag for protein purification. Cloning details are available upon request.

### Recombinant protein production

Sf9 cultures were grown in SF900 media and infected with Cdc13-encoding baculovirus at a multiplicity of infection of 0.1. Cells were harvested by centrifugation after 3 days at 27°C with gentle aeration, and pellets were stored at -80°C. Frozen cell pellets were lysed in the presence of protease inhibitor cocktail (aprotinin, chymostatin, leupeptin, and pepstatin A at 5 μg/mL and 1 mM phenyl-methyl-sulfonyl fluoride), as well as 10 *μ*g/mL DNase I, and clarified lysates were purified using nickel resin followed by anti-FLAG resin as previously reported (9). Briefly, cells were disrupted by sonication for 3 min (15 s sonication, 30 s rest; 12 times), and lysates were clarified by ultracentrifugation (20,000 x g for 20 min) and incubated with 0.4 mL of His-Select nickel resin (Sigma-Aldrich) for 1 h at 4°C with gentle mixing. The resin was pelleted at 1000 g for 5 min and transferred to a gravity column. The matrix was washed twice with 10 mL K buffer containing 500 mM KCl, 0.1% NP-40, and 15 mM imidazole and then washed once with 10 mL of K buffer containing 500 mM KCl, 0.01% NP-40, and 15 mM imidazole. Protein was eluted with 2.5 mL of K buffer containing 200 mM imidazole. The eluate was then gently mixed with 300 μL of anti-FLAG-M2 resin (Sigma-Aldrich), incubated for 1.5 h or overnight at 4°C, and washed as above. The target protein was eluted twice by incubating the matrix with 0.5 mL of K buffer supplemented with 200 μg/mL FLAG peptide (Sigma) for 0.5 h, followed by a third elution step with no incubation. The eluate was filter-dialyzed into K buffer in an Ultracel-30K concentrator (Millipore, Burlington, MA, USA) and further concentrated to ∼100 μL. Purity was assessed by SDS-PAGE and western blotting analyses. Purified His_6_-Cdc13-FLAG protein was divided into 5-μL aliquots, frozen in liquid nitrogen, and stored at -80°C. Cdc13-L91R was purified identically to wild-type Cdc13. All protein preparations were assessed for DNA binding activity and contaminating nuclease activity using standard gel shift assays prior to experimental use.

### Mass photometer analysis

Experiments were performed using a Two^MP^ mass photometer (Refeyn, Oxford, UK) to determine the oligomeric state of recombinant Cdc13 and Cdc13-L91R in solution. The protein mass standards BSA (66.5, 133, and 199.5 kDa for monomers, dimers, and timers, respectively) and thyroglobulin (670 kDa) were used to generate a calibration curve for molecular weight determinations. All mass photometer experiments were performed in 1x DNA binding buffer (25 mM HEPES (pH 8.0), 50 mM NaOAc, 150 mM NaCl, and 7.5 mM MgCl_2_) at room temperature. Proteins were diluted to 15 nM and incubated for 30 min at 30°C in the presence or absence of 50 nM ssDNA substrate. The mass photometer was focused with 1x binding buffer, and proteins and protein-DNA complexes were diluted on the sample slides to 11.25 nM protein. Samples were analysed following the manufacturer’s protocols, and the data were graphed using DiscoverMP software.

### DDE assay

The DDE assays were performed as described (9). Briefly, IR800-labeled telomeric oligonucleotide substrate was mixed to a final concentration of 2 nM on ice with 3.75 nM wild-type Cdc13 (dimer concentration) or Cdc13-L91R (monomer concentration). All binding reactions were performed in a volume of 9 µL and were incubated for 15 min at 30°C in 1x binding buffer containing 5% (w/v) glycerol and 0.01% Tween-20 (w/v). These conditions led to saturated binding for both proteins. Competitor ssDNA conjugated to an IR700 label was then titrated into the binding reactions and incubated for an additional 15 min at 30°C. Reaction products were separated on 8% native acrylamide gels (37.5:1 acrylamide:bisacrylamide) run in 1x Tris-glycine running buffer (25 mM Tris and 185 mM glycine (pH 8.8)) at 100 V for 30-45 min. Gels were visualized using a LI-COR scanner and images were quantified using Image Studio Lite software, version 5.2.

### BLI

BLI experiments were performed to determine association, dissociation, and DDE rates using a BLItz instrument (FortéBio, Fremont, CA, USA) following the manufacturer’s protocols. All binding assays were performed in 1x DNA binding buffer at room temperature (∼23°C). Streptavidin sensors were saturated by binding to 1 µM biotinylated Tel30G for 2 min. Sensors were then moved to a 100 nM solution of either wild-type Cdc13 or Cdc13-L91R. Association of the protein to immobilized DNA was recorded for 2 min. Then, sensors were shifted to a large volume (600 µL) of 1x binding buffer, and dissociation was recorded for an additional 2 min. Finally, the BLI sensor was moved to a new solution with 10 nM unlabelled Tel30G to observe DDE, and data were recorded for 2 more minutes. The rates of association, dissociation, and DDE were calculated using GraphPad Prism software.

### Statistical analyses

Biological replicates (≥3) were performed for all assays. Unless otherwise noted, the plotted data points are the means of these replicates, and the error bars are the standard deviation. All graphs were plotted using GraphPad Prism software, and where curves are fitted to the data, the details about curve fitting are provided in the figure legends.

### Structure prediction

AlphaFold 3 models were rendered with both protein and DNA on the Google DeepMind AlphaFold Server (alphafoldserver.com) using default parameters unless otherwise noted. The protein sequence for *S. cerevisiae* Cdc13 from the S288c genetic background was obtained from the *Saccharomyces* Genome Database (yeastgenome.org). The sequences for telomeric ssDNA and a telomere-like substrates (*i*.*e*., duplex DNA with a 3⍰ G-strand ssDNA tail) were input *in silico* with K^+^ ions. See Table S1 for additional details. Structures obtained from AlphaFold 3 were visualized and analyzed in Chimera X (http://www.cgl.ucsf.edu/chimerax).

## RESULTS AND DISCUSSION

Having previously characterized the nucleic acid characteristics and Cdc13 domains required for DDE, we here sought to determine if Cdc13 dimerization is also required. However, we first needed to generate a dimerization-deficient mutant. Cdc13 behaves as an obligate homodimer both *in vivo*, by the criteria of co-immunoprecipitation and yeast two-hybrid assays, and *in vitro*, as demonstrated with recombinant protein and chemical crosslinking and sucrose gradient sedimentation assays (22). It has been reported that the OB1 domain mediates this dimerization (22,40) and that L91A (40) and L91R (22) mutations disrupt it in the context of isolated recombinant OB1 domain preparations. In the context of the full-length Cdc13 protein, SEC-MALS analysis indicates that the L91R mutation also disrupts dimerization, though this mutant protein still displays positive cooperativity when binding to ssDNA (8), suggesting that the substrate may induce dimerization. Further, AlphaFold 2 structural prediction of Cdc13 homodimers indicates that the OB2 and OB4 domains also mediate intra- and inter-monomer contacts (9), calling into question the strength of the L91R effect.

### The Cdc13-L91R mutation disrupts dimerization in solution

To unambiguously determine if Cdc13-L91R is monomeric in solution, both in an apo state and bound to ssDNA, we generated recombinant protein and analysed it by mass photometry. We typically dilute protein preparations using PBS for mass photometry analysis but found that these buffer conditions alone disrupt wild-type Cdc13 homodimers, yielding a mixed population of monomers (∼105 kDa) and dimers (∼210 kDa) in solution (Fig. 1A). Pre-binding Cdc13 to a 30-nt telomeric repeat sequence ssDNA substrate (Tel30G) prior to PBS dilution yielded a single mass peak with a molecular weight consistent with a dimer of Cdc13 bound to an approximately 10-kDa ssDNA (Fig. 1B; (42)). The analysis does not preclude the possibility of two Cdc13 monomers independently binding the same substrate, but one would expect to observe both one-monomer and two-monomer binding events if that were the case. Regardless, when the mass photometry analysis was conducted using the same buffer that is used in the ssDNA binding assays, only Cdc13 dimers were observed (Fig. 1C). Similarly, when only using binding buffer, both single-dimer and double-dimer binding was observed upon the addition of an even longer 50-nt telomeric repeat sequence ssDNA substrate (Tel50G; ∼17 kDa) that can accommodate two Cdc13 dimers (Fig. 1D). Thus, the Cdc13-ssDNA binding buffer was used for all subsequent mass photometry experiments.

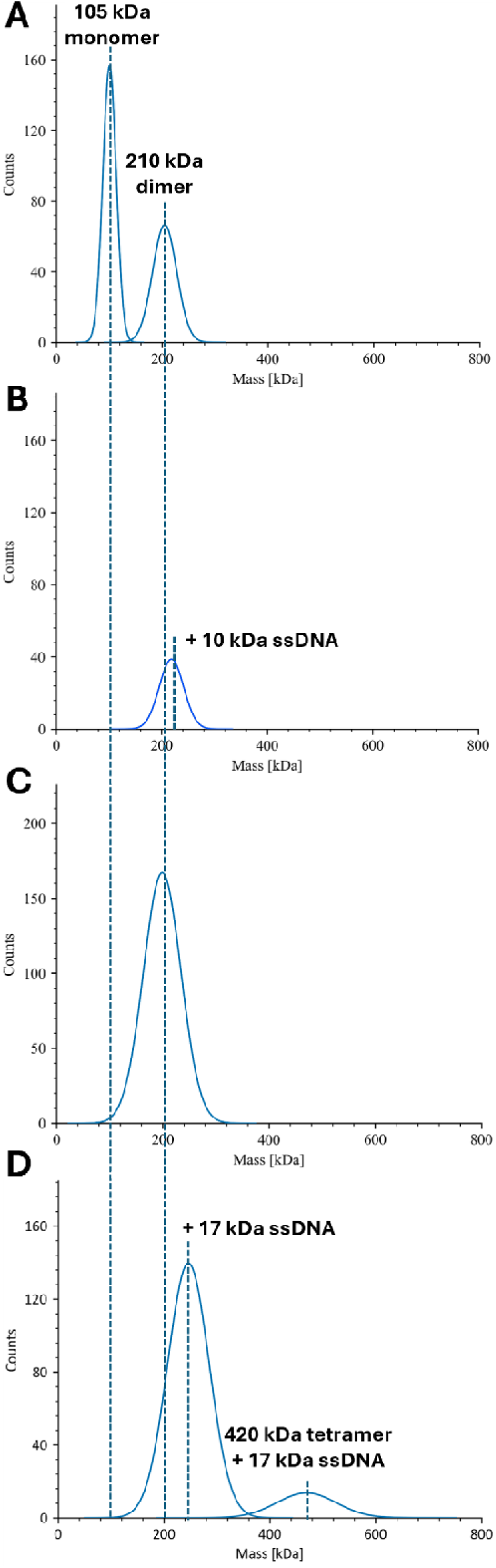

When analysing recombinant Cdc13-L91R, we found that it only formed monomers in solution (Fig. 2A), consistent with the previously published SEC-MALS work (8). Binding to Tel30G also only yielded a single mass peak consistent with a monomer bound to one strand of ssDNA (Fig. 2B), confirming that ssDNA binding does not induce Cdc13-L91R dimerization. However, binding to the longer Tel50 ssDNA did produce a small population of molecules with a molecular mass consistent with two Cdc13-L91R proteins (Fig. 2C), though we interpret these to be independent protein molecules bound to the same substrate rather than a protein dimer. In summary, our recombinant Cdc13 wild-type and L91R protein preparations exist as homodimers and monomers, respectively, in solution in the buffer conditions used to analyse DDE, so these reagents can conveniently be used to determine if dimerization is necessary for DDE.

**Figure 2.**
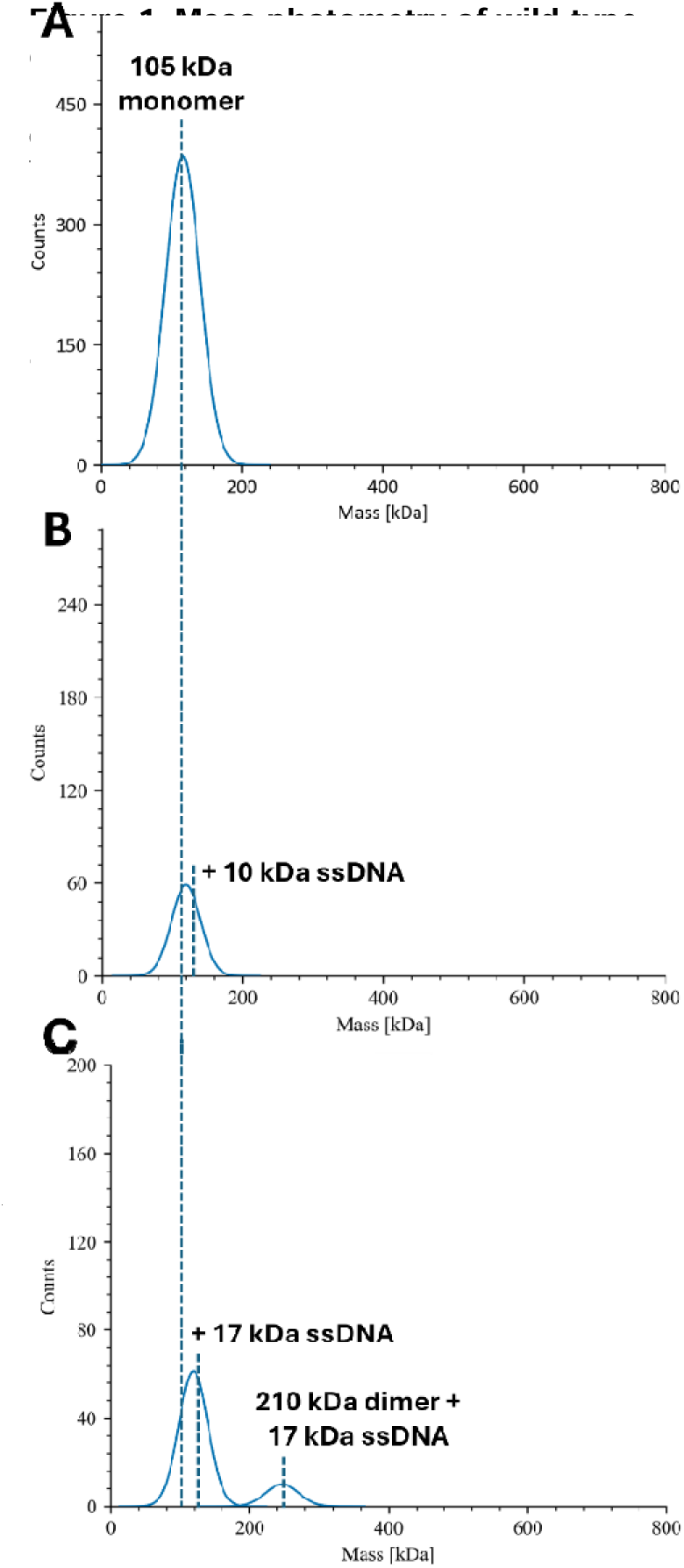
Cdc13-L91R is monomeric is solution. A) Recombinant Cdc13-L91R exists as a monomers in solution, and ssDNA binding (Tel30G) does not induce dimerization (B). C) Tel50G can support the binding of two Cdc13-L91R monomers. All assays contained 11.25 nM protei**n** and 50 nM ssDNA where indicated.

### Cdc13 dimerization is necessary for DDE

To assay for DDE, we typically use a Tel30G sequence conjugated to one of two different near-infrared fluorophores: IR700 and IR800 (9). By prebinding Cdc13 to saturating amounts of IR800-Tel30G and then titrating in IR700-Tel30G (or *vice versa*), we can monitor the exchange of the protein from one substrate to the other using a gel-based assay and two-colour fluorescent imaging. We define DDE as sub-stoichiometric concentrations of secondary substrate being able to effectively compete for Cdc13 binding. In contrast, a large molar excess of secondary substrate is necessary for typical competition binding, which is driven by the off-rate of the protein for its substrate.

An example of wild-type Cdc13 undergoing DDE is shown in Figure 3A, where Cdc13 binding to IR800-Tel30G is disrupted by sub-stoichiometric concentrations of IR700-Tel30G. In contrast, much higher concentrations of IR700-Tel30G are required to compete for Cdc13-L91R binding (Fig. 3B). These assays were performed in biological triplicates, and the results are quantified in Figure 3C. As reported previously (9), the EC_50_ of Cdc13 bound to IR800-Tel30G and challenged with IR700-Tel30G as a secondary substrate is 0.89 nM, but that of Cdc13-L91R is 31.6 nM, a 35.5-fold difference. Because it took a large stoichiometric excess of secondary substrate to compete off Cdc13-L91R, we concluded that Cdc13-L91R is defective for DDE, and thus, DDE requires Cdc13 dimerization.

**Figure 3.**
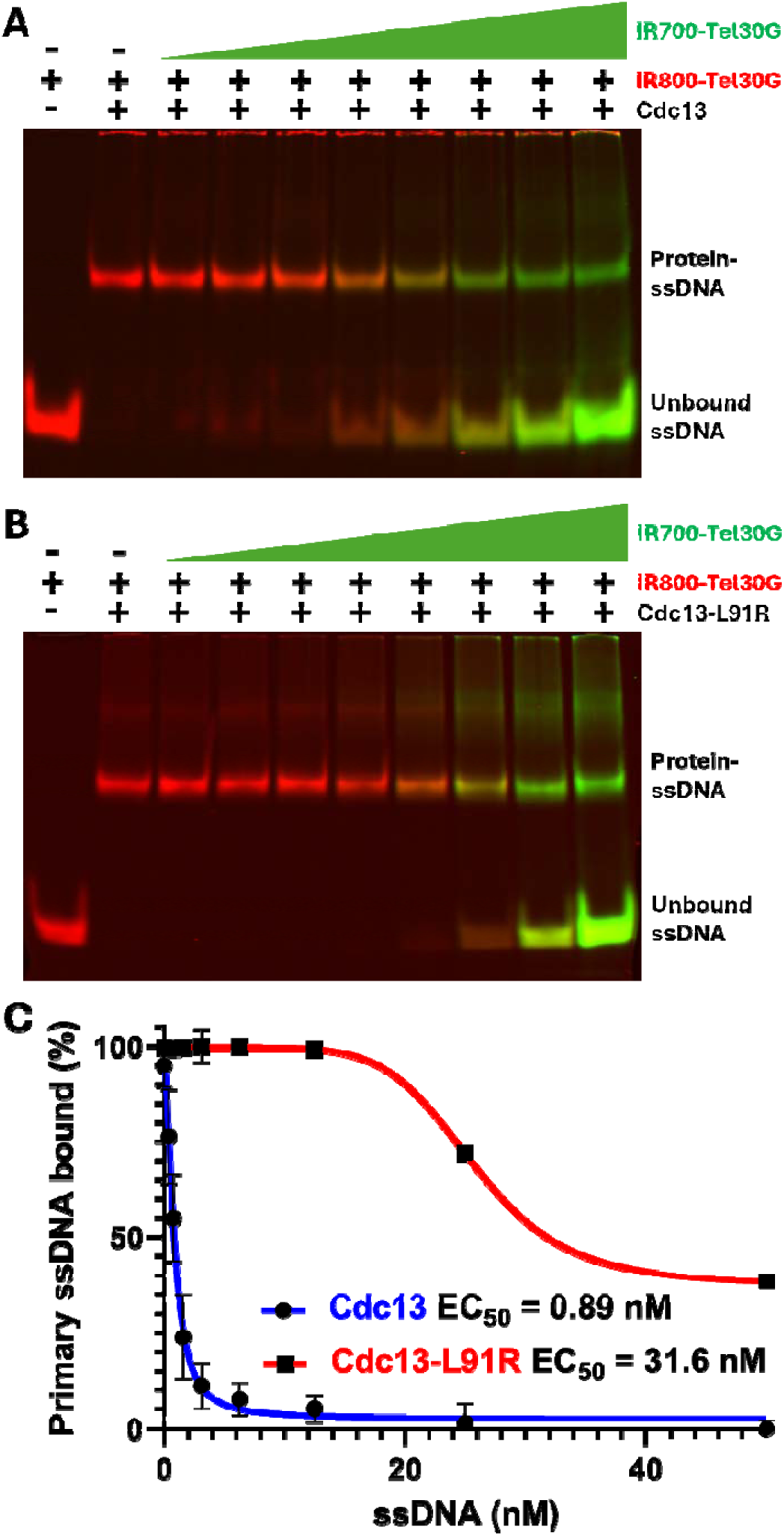
Dual-label dissociation assays demonstrate that Cdc13-L91R lacks DDE activity. A) Representative gel image of the dissociation of 3.75 nM wild-type Cdc13 dime**r** prebound to 2 nM IR800-Tel30G and exposed to increasing concentrations (0.2-15 nM) of IR700-Tel30G. B) Representative gel image of the dissociation of 3.75 **nM** Cdc13-L91R monomer prebound to 2 nM IR800-Tel30G and exposed to increasing concentrations of IR700-Tel30G. C) Graph of the displacement of the initially bound IR700-Tel30G DNA with Cdc13 wt or Cdc13-L91R. Error bars represent the

### Cdc13-L91R does not undergo DDE

An orthogonal method to observe DDE in real time is by using BLI (9). Cdc13-ssDNA complexes are reported to be extremely stable, with a half-life approaching 2 days (23), while DDE occurs on the time scale of seconds (9). Both of these phenomena can be observed for wild-type Cdc13 by BLI, where its off-rate from a Tel30G ssDNA is miniscule over at least 24 h unless a secondary pool of Tel30G substrate is introduced to initiate DDE (9). Here, we used BLI to recapitulate these Cdc13-Tel30G binding results and to further characterize Cdc13-L91R (Fig. S1).

Both wild-type and L91R Cdc13 preparations bind Tel30G with similar association kinetics (Fig. 4A). However, they differ in the pre-DDE dissociation phase of the assay (Fig. 4B). As reported (9), full-length wild-type Cdc13 undergoes a brief initial dissociation and then reassociation with the ssDNA upon dilution of the protein-DNA complexes into a large volume of buffer, but then no further dissociation is observed. We previously hypothesized that this could be due to local dissociation and reassociation of Cdc13 with the ssDNA substrates on the molecularly crowded BLI sensor tip. Alternatively, it could be due to condensation and then relaxation of the ssDNA conjugated to the sensor because Cdc13 condensation of immobilized DNA has been reported (35). Regardless, Cdc13 truncation mutants that do not undergo DDE lack this phase in their BLI profiles (9), and likewise, Cdc13-L91R lacks it here (Fig. 4B). However, unlike Cdc13 truncation mutants, which all display similar levels of protein dissociation from the ssDNA (9), Cdc13-L91R actually appears to continue to slowly associate with the Tel30G on the sensor. It is not until secondary substrate is flooded into the reaction during the DDE/competition step that Cdc13-L91R begins to dissociate from the ssDNA (Fig. 4C). It experiences a rapid initial drop in binding, followed by a slow and steady decrease. Although wild-type Cdc13 likewise undergoes an initial rapid drop in binding, its subsequent dissociation is more rapid than Cdc13-L91R, consistent with the protein undergoing wild-type dimeric Cdc13 DDE as opposed to the simple binding competition displayed by monomeric Cdc13-L91R. This lack of DDE along ssDNA could explain why shortened telomeres are observed *in vivo* with the Cdc13-L91R mutant (40).

**Figure 4.**
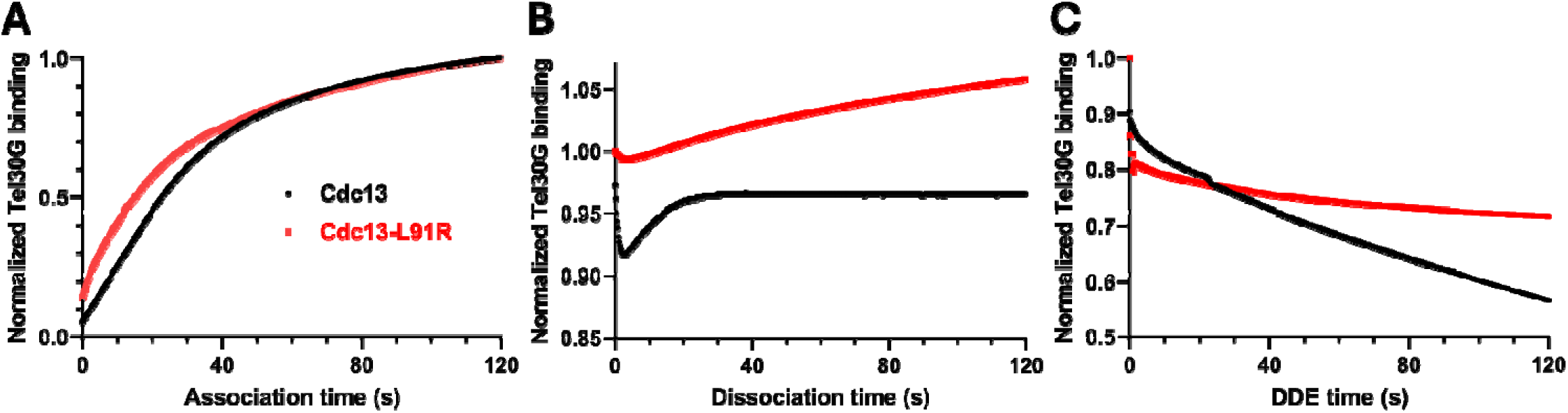
BLI analysis of Cdc13 and Cdc13-L91R binding to telomeric ssDNA. A) The association of 100 nM wild-type Cdc13 or Cdc13-L91R with immobilized Tel30G ssDNA was monitored for 2 min. B) Then, the protein-ssDNA complexes were diluted into a large volume of DNA-free buffer, and dissociation of the proteins from Tel30G was monitored for 2 min. C) Finally, the remaining protein-ssDNA complexes were exposed to free unlabelled Tel30G ssDNA in solution, driving either DDE (Cdc13) or simple binding competition (Cdc13-L91R). The curves shown in all plots are comprised of data points collected every 0.2 s and are representative of ≥ 3 independent experiments. The wild-type Cdc13 data are from (9) and were collected at the same time as the Cdc13-L91R dat .

Failure of Cdc13 to dimerize could also interfere with DNA polymerase α-primase or Est1 interactions that would lead to shortened telomeres (13,22,40).

### Models for DDE based on the Cdc13 dimer

The results described above raise interesting questions about the mechanism of DDE by Cdc13 and the role dimerization might play in this process. Although our *in vitro* assays only monitor intermolecular DDE between two different ssDNAs, we hypothesize that intramolecular DDE from one binding site to another along a single long telomeric ssDNA is possible. This notion is based in part upon similar phenomena displayed by ssBPs, such as the long-range lateral diffusion demonstrated by RPA in single-molecule experiments (43). It is likewise based on the fact that Cdc13 must transition from the dsDNA-ssDNA junction at a short telomere toward the 31.-end of a growing telomere to support telomerase activity and fill-in by Pol *α*-primase (11-17). Crucially, there is no known mechanism for Cdc13 movement, but intramolecular DDE could explain it.

The results from our mass photometer studies showing that Cdc13-L91R is a monomer in solution (Fig. 2), combined with the lack of DDE activity displayed by the L91R mutant relative to wild-type Cdc13 (Fig. 3), leads us to propose that dimerization of Cdc13 is critical for its ability to move along telomeric DNA. Given that there is one high-affinity and one low-affinity ssDNA binding domain per monomer of Cdc13, the L91R results imply that ssDNA binding across monomers or allosteric communication of the ssDNA bound state of one monomer to another within the dimer, is crucial for DDE activity (Fig. 5). Because Cdc13 binding to the long Tel50G substrate significantly inhibits DDE activity, this suggests the possibility that telomeric ssDNA ≥ 50 nt is bound in a manner that fills all effective Cdc13 ssDNA binding sites. Based on our model in Figure 5, this could mean filling both OB1 and OB3 domains in the dimer (four sites filled) or both high-affinity sites (OB3s) and one low-affinity site (OB1 in subunit A or B). Binding to ssDNA < 50 nt long (*e*.*g*., Tel30G) leaves some ssDNA binding domains open for additional interaction by Cdc13 and potential movement along telomeric DNA.

**Figure 5.**
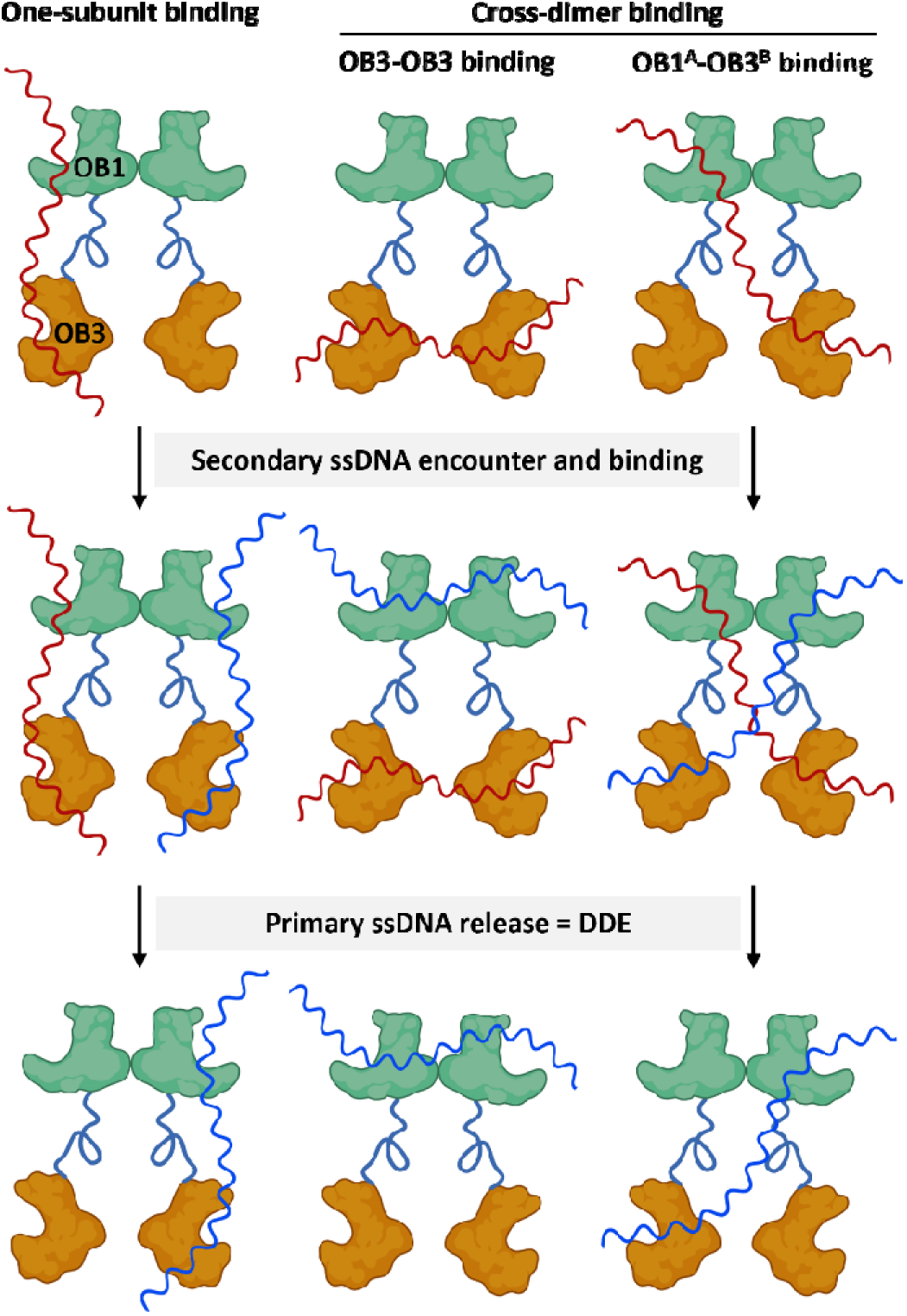
Cdc13 dimer models for DDE. There are four known ssDNA binding sites in a Cdc13 homodimer, *i*.***e***., one OB1 domain and one OB3 domain from each subunit. A long enough ssDNA substrate (top, red) can fill both OB1 and OB3 sites in one monomer (left), fill both high-affinity OB3 sites across the dimer (middle), or span the OB1 domain **of** subunit ‘A’ in the dimer and the OB3 domain of subunit ‘B’ (or *vice versa*). All three scenarios leave two open ssDNA binding sites that can be filled when encountering a secondary ssDNA substrate (blue, middle). Upon filling the second set of binding sites, the binding event is allosterically communicate**d** across the dimer, perhaps via the OB2 and/or OB4 domains (not shown), leading to release of the primary ssDNA to complete the dynamic exchange of substrates (DDE, bottom). These models are depicted with two separate ssDNAs, but **a** single substrate of sufficient length, especially one that is actively being lengthened by telomerase, is predicted to fulfill the same role.

### Cdc13-ssDNA complex structure prediction

As a first step to understanding how Cdc13 binding to telomeric ssDNA of increasing length might regulate DDE and/or conformational changes in Cdc13 homodimers, we turned to protein structure prediction of Cdc13 in the presence and absence of telomeric DNA. Here, AlphaFold 3 predicts that the major Cdc13 dimer interface is the OB1 domain in each monomer when no ssDNA is present (Fig. 6A). This corresponds to the published work (22,40) and that presented here on the Cdc13-L91R mutation ablating dimerization (Fig. 2), but it differs from the AlphaFold 2 model that we previously reported, in which the OB2 and OB4 domains also participate in dimerization (9). That said, weak inter-subunit contacts are formed between the OB2 and OB4 domains when the Cdc13 dimer is modelled bound to telomeric ssDNA (Fig. 6B-E), and OB1-OB1 and OB1-OB3 interactions are strengthened compared to apo-Cdc13 (Fig. 6A). As the length of telomeric ssDNA bound to Cdc13 increases from 11 to 50 nt, the conformation of the protein also progressively changes (Fig. 6A-D). Conformational changes are also evident when Cdc13 is modelled bound to one telomere-like substrate (*i*.*e*., dsDNA with a 3’ ssDNA tail) *vs*. two telomere-like substrates (Fig. 7). Interestingly, Figure 7B shows that the Cdc13 dimer is predicted to synapse two telomeres, which would support the cross-dimer ssDNA binding model in Figure 5. Although AlphaFold structural predictions should be interpreted with caution, we do note that the predicted structures of the OB1 and OB3 domains in the Tel30G-bound model can be superimposed showing that they have similar structures (Fig. S2), as was previously demonstrated using actual X-ray crystal structure data of these domains (22).

**Figure 6.**
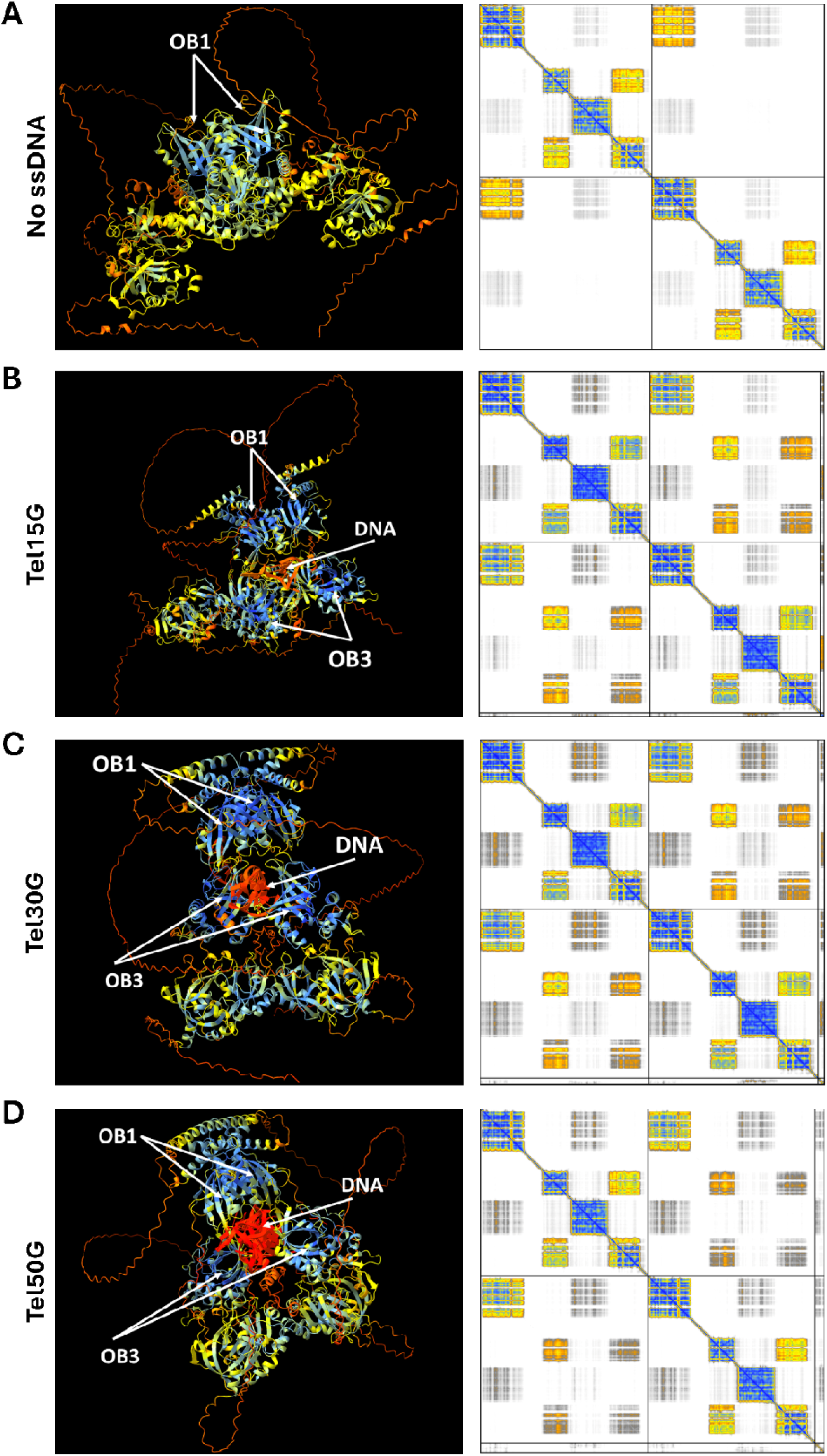
AlphaFold modelling of apo- and ssDNA-bound Cdc13 dimers. The AlphaFold 3 server was used to model the Cdc13 dimer (A) and the dimer bound to 15 (B), 30 (C), or 50 nt (D) of telomeric repeat sequence ssDNA. The positions of the OB1 and OB3 domains, as well as the ssDNA substrate, are labelled. Predi**c**ted alignment error plots are shown to the right of each structure. Intra-subunit contacts are shown in the top left and bottom right squares; inter-subunit contacts ar**e** shown in the others. Protein-ssDNA contacts are Cooler colors indicate higher confidence in the prediction.

**Figure 7.**
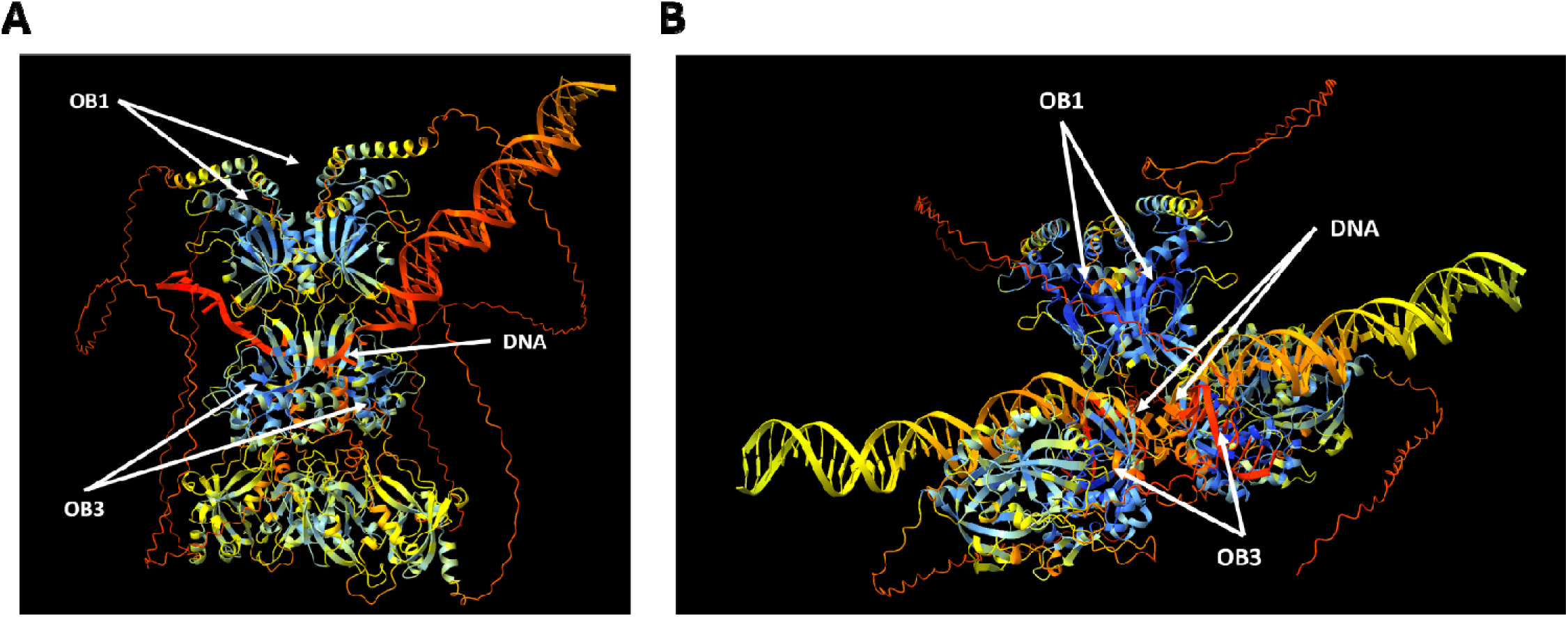
Predicted structures of Cdc13 homodimers bound to one (A) and two (B) telomeres. The telomere-bound Cdc13 dimer structures were predicted using the AlphaFold 3 server.

Combining our results with published protein structural data and the AlphaFold 3 predictions, we propose a refined model of Cdc13 binding and movement along telomeric DNA. As DNA replication is nearing completion, Cdc13 interacts with DNA polymerase α to transition from telomerase interaction to fill-in of the lagging (C-rich) strand. On leading strands, resection of the C-rich strand is necessary to create the 3□ overhang that is the Cdc13 binding site. This leaves telomeres with 12-18 nt of 3□ ssDNA protected by the CST complex. As the next replication cycle begins, Stn1 and Ten1 dissociate from Cdc13, allowing the Cdc13 telomerase recruitment domain to interact with the Est1 subunit of telomerase. If active telomere extension occurs, our model suggests that Cdc13 will dynamically exchange with newly synthesized telomere DNA at a rate of exchange proportional to the length of new telomeric ssDNA. This movement would be limited to the 3□ direction due to dsDNA blocking movement in the 5□ direction. This Cdc13 movement will continue until the size of the ssDNA G-strand reaches ≥ 50 nt, which significantly reduces the rate of DDE activity. As this occurs, the Cdc13 conformation shifts in a manner that favours binding of Cdc13 to other interaction partners such as DNA polymerase α and Stn1/Ten1. Finally, our data demonstrates that DDE activity requires both OB1 and OB3 domains, as well as homodimers of Cdc13. Cdc13 may bind ssDNA with a single OB3 domain, both OB3s, or utilize a mixed OB1-OB3 binding site. When additional telomeric ssDNA has been synthesized of sufficient length, it can be bound by the open, unoccupied Cdc13 ssDNA binding site(s), communicating the status of this secondary binding, and releasing the initially bound ssDNA. The precise mechanism for DDE inhibition by long (∼50 nt) ssDNAs is unknown, but it may involve complete filling of all potential Cdc13 ssDNA binding domains, stabilizing the protein-DNA interaction.

## Conclusions

George Box famously wrote that, “all models are wrong,” but also stated that “The good scientist must have the flexibility and courage to seek out, recognize, and exploit such errors – especially his own” (44). In other words, all models are wrong, but some are useful. It is our hope that the models presented above fall into the latter category, but as with AlphaFold structural predictions, they should also be interpreted with caution. For instance, it is clear that Cdc13 functions both as a homodimer and as part of the CST complex *in vivo*, but the stoichiometry of the *S. cerevisiae* CST complex is unknown. Are two copies of each protein present in the complex, such that the Cdc13 homodimer is maintained? If so, can CST also undergo DDE, or does the interaction with Stn1 and Ten1 inhibit DDE activity? The analogous human complex (CTC1-STN1-TEN1) displays 1:1:1 stoichiometry in solution, with 10 CST complexes assembling into a ring-shaped super-complex when bound to (TTAGGG)_3_ ssDNA, as demonstrated by cryo-electron microscopy (45). However, at the primary sequence and three-dimensional structure levels, CTC1 shares little homology with Cdc13. Further complicating this issue is the fact that the more closely related CST complex from the yeast *Candida glabrata* di plays a 2:4:2 or 2:6:2 stoichiometry between its Cdc13, Stn1, and Ten1 subunits (46), though direct, high-resolution analyses of the *C. glabrata* CST complex remain to be performed. Thus, answering our questions will require the generation of recombinant *S. cerevisiae* CST complexes that are stable in solution and competent for ssDNA binding for analysis as described above and in (9).

Similarly, all of our *in vitro* DDE data are based on intermolecular exchange of Cdc13 between substrates, while many of our hypotheses consider intramolecular Cdc13 movement on one telomeric ssDNA G-tail as it is being elongated. Inter-telomere ssDNA binding and/or DDE by Cdc13 could serve as a means to support telomere biology, for instance aiding in telomere bouquet formation during meiotic prophase (4) or providing a means for a single Cdc13 dimer to act at multiple telomeres during a single cell cycle. Indeed, the latter may be necessary because Cdc13 is a low-abundance protein (47-49) with a high abundance of potential binding sites throughout the yeast genome. Regardless, testing for intramolecular DDE of Cdc13 on ssDNA will require single-molecule techniques, such as those used to monitor movement of RPA on ssDNA (43). Our models in Figure 5 also invoke the use of all four ssDNA binding sites in a Cdc13 homodimer, but the mutants tested to date either impact both subunits in the dimer or yield monomers. Thus, determining the mechanism of DDE by Cdc13 will require the generation of mixed dimers containing one or more ssDNA binding-deficient domains in the context of otherwise wild-type OB folds. Mixed dimers of differentially tagged Cdc13 have been observed *in vivo* (22), so a tandem affinity purification scheme for differentially tagged recombinant Cdc13 and Cdc13 mutants could yield sufficient material to test the models in Figure 5.

In summary, we were the first to observe and report DDE by Cdc13, and we demonstrate here that Cdc13 homodimerization is necessary to support this activity. The physiological implications of DDE have yet to be investigated, but it is our hope that the models, hypotheses, and unanswered questions highlighted above will guide further *in vivo* and *in vitro* investigation of this phenomenon.

## DATA AVAILABILITY

The data underlying this article are available in the article, in its online supplementary material, or will be shared upon reasonable request to the corresponding author.

## SUPPLEMENTARY DATA

Supplementary Data are available at NAR online.

## ACKNOWLEDGEMENTS

We thank Hengyao Niu and members of his laboratory for sharing plasmids and for helpful advice with insect cell culture and protein purification.

## AUTHOR CONTRIBUTIONS

David Nickens: Conceptualization, Formal analysis, Methodology, Validation, Writing—original draft. Spencer Gray: AlphaFold modeling, Formal analysis, Methodology, Writing—review & editing. Robert Simmons: Mass photometry, Writing—review & editing. Matthew Bochman: Conceptualization, Formal analysis, Methodology, Validation, Writing— review & editing.

## SUPPLEMENTARY MATERIALS

### Supplementary Figures

**Figure S1..**
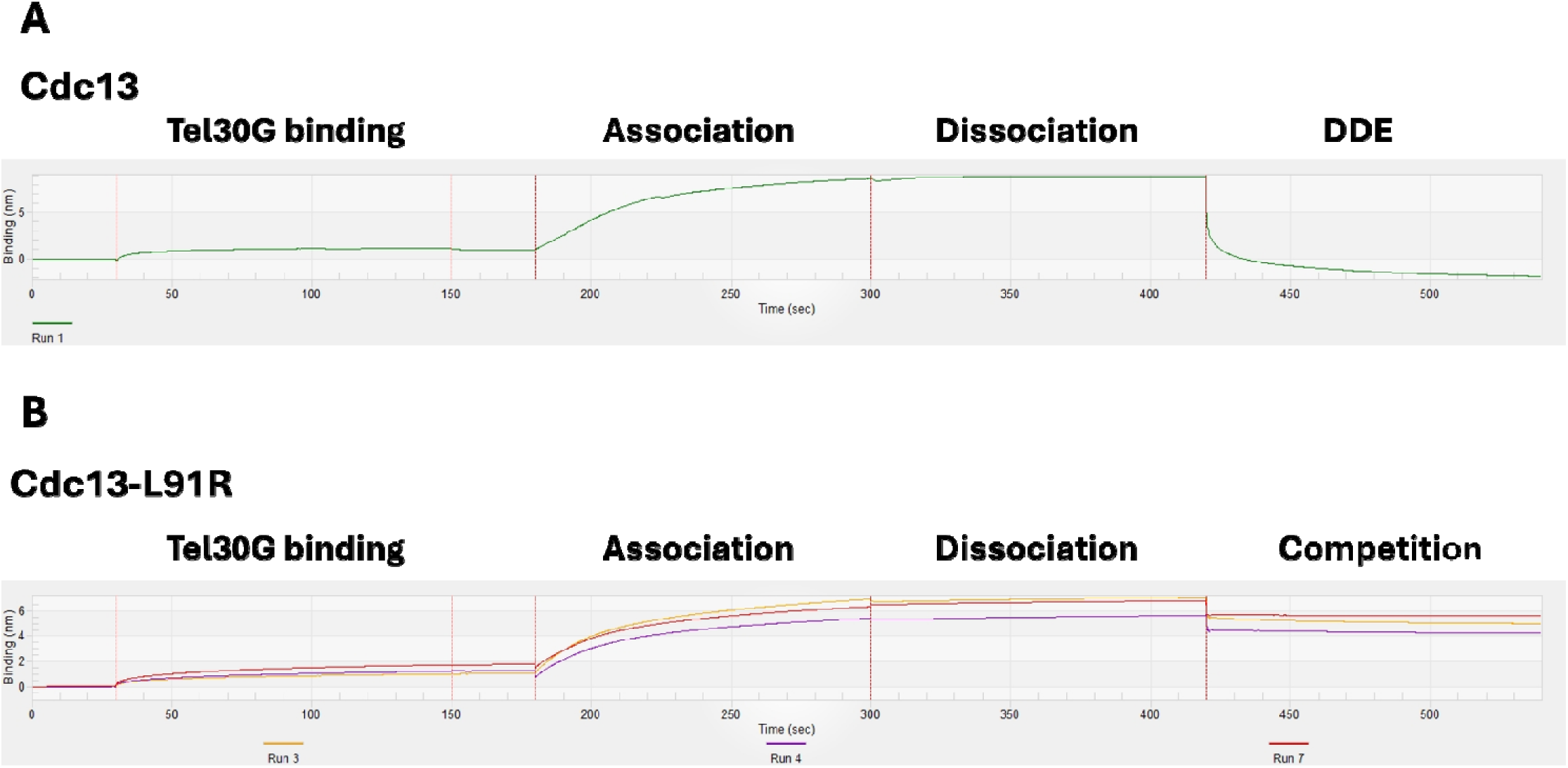
Example BLI assays to observe DDE. In a typical assay, biotinylated Tel30G ssDNA was immobilized on a streptavidin-coated BLI sensor (Tel30G binding), washed, exposed to recombinant protein for ssDNA binding (Association), exposed to a large volume of buffer to allow dissociation of the protein from the ssDNA (Dissociation), and then exposed to a large volume of buffer containing a molar excess of competitor ssDNA to allow DDE or binding competition to occur (DDE/Competition).

**Figure S2..**
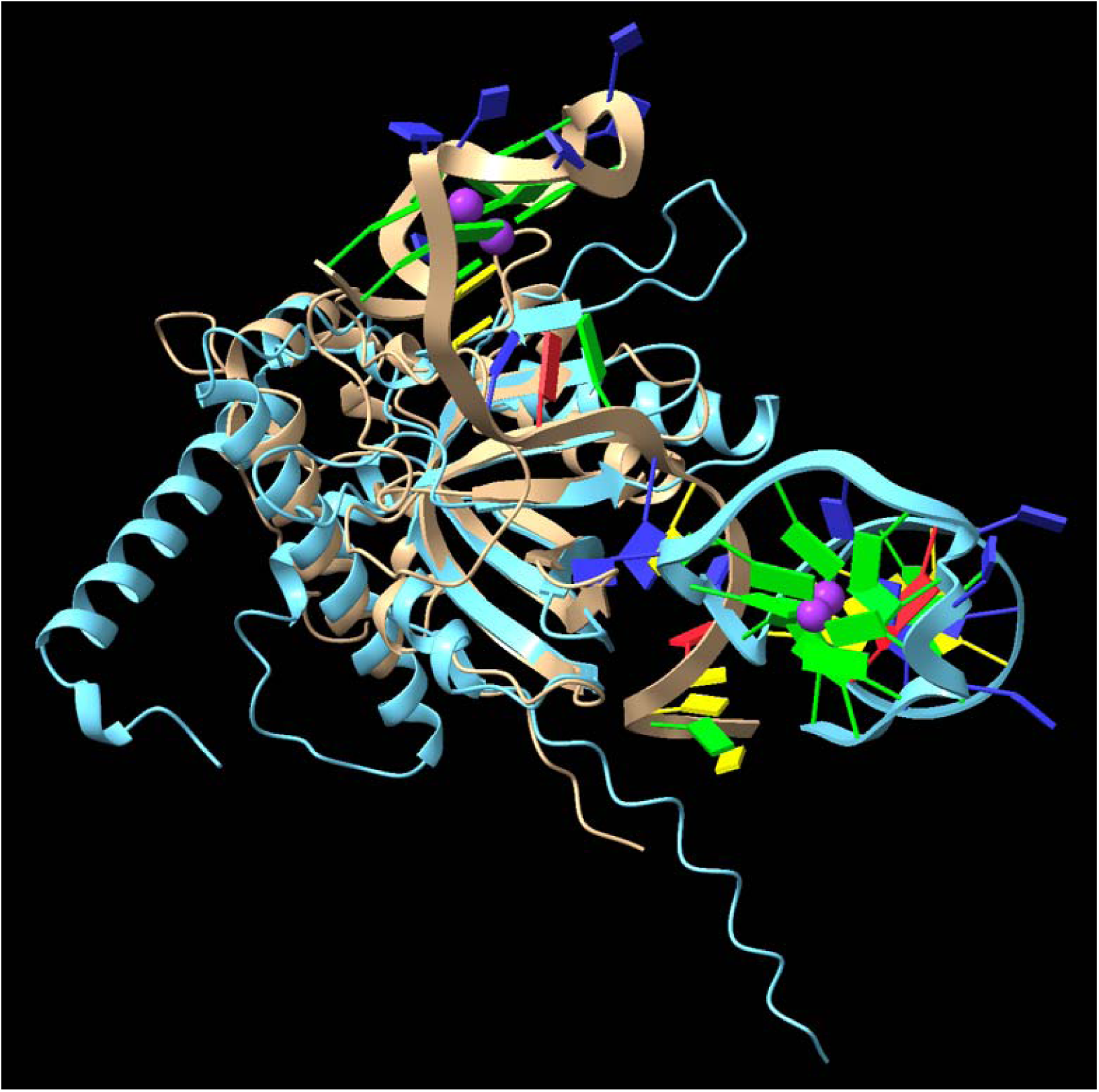
The OB1 domain (light blue) of Cdc13 was superimposed with the Cdc13 OB3 domain (tan) bound to Tel30G. Both structures were obtained from AlphaFold 3 prediction. See Table S1 for the domain boundaries used relative to full-length Cdc13.

### Supplementary Tables

**Table S1.**
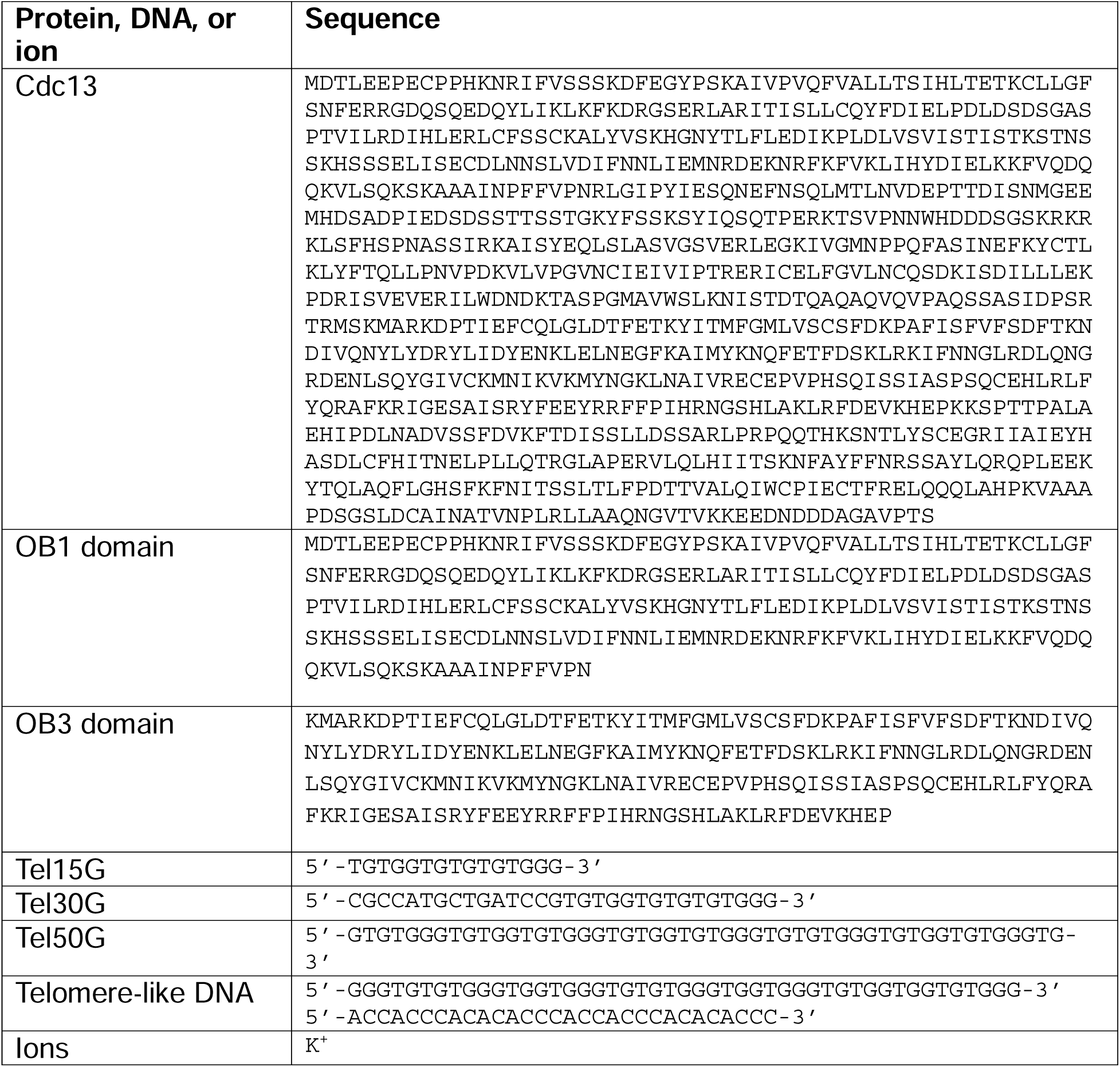
AlphaFold 3 inputs.

